# GAPDH cooperativity mediates drug resistance and metabolism in *Plasmodium fal-ciparum* malaria parasites

**DOI:** 10.1101/2021.11.05.467329

**Authors:** Andrew J. Jezewski, Ann M. Guggisberg, Dana M. Hodge, Naomi Ghebremichael, Lisa K. McLellan, Audrey R. Odom John

## Abstract

Efforts to control the global malaria health crisis are undermined by antimalarial resistance. Identifying mechanisms of resistance will uncover the underlying biology of the *Plasmodium falciparum* malaria parasites that allow evasion of our most promising therapeutics and may reveal new drug targets. We utilized fosmidomycin (FSM) as a chemical inhibitor of plastidial isoprenoid biosynthesis through the methylerythritol phosphate (MEP) pathway. We have thus identified an unusual metabolic regulation scheme in the malaria parasite through the essential glycolytic enzyme, glyceraldehyde 3-phosphate dehydrogenase (GAPDH). Two parallel genetic screens converged on independent but functionally analogous resistance alleles in GAPDH. Metabolic profiling of FSM-resistant *gapdh* mutant parasites indicates that neither of these mutations disrupt overall glycolytic output. While FSM-resistant GAPDH variant proteins are catalytically active, they have reduced assembly into the homotetrameric state favored by wild-type GAPDH. Disrupted oligomerization of FSM-resistant GAPDH variant proteins is accompanied by altered enzymatic cooperativity and reduced susceptibility to inhibition by free heme. Together, our data identifies a new genetic biomarker of FSM-resistance and reveals the central role of GAPDH cooperativity in MEP pathway control and antimalarial sensitivity.

## INTRODUCTION

Worldwide, malaria causes more than 400,000 fatalities each year, predominantly in young infants and pregnant women (1). Successful clinical management of life-threatening blood-stage infections with *Plasmodium falciparum* relies on a rapid response to antimalarial chemotherapeutics (2, 3). Because of widespread antimalarial resistance to older agents, the newer artemisinin-based combination therapies (ACTs), which quickly reduce parasite loads, are first-line therapeutic agents (4). Expanding resistance to ACTs, characterized by delayed parasite clearance during treatment, has created a pressing public health need to develop antimalarials with novel modes of action (5–8). Identifying novel biological processes that may serve as useful drug targets is an important approach in the ongoing efforts toward malaria elimination (9, 10). A deep understanding of essential parasite biology, particularly those parasite processes that are distinct from the human host, will open new possibilities for therapies effective against drug-resistant *P. falciparum.*

*P. falciparum* possesses an unusual plastid organelle termed the apicoplast, which was originally acquired through an ancient endosymbiotic event of a plastid-bearing red alga (11, 12). While photosynthetic capacity has been lost over evolutionary time, this unique organelle has been retained by most modern Apicomplexans because it contributes essential metabolic products (13, 14). In particular, isoprenoid precursors generated by the apicoplast are essential for asexual development of *P. falciparum* (15, 16). Isoprenoids are a diverse set of biomolecules that contribute to organismal color (carotenoids) and odor (volatiles), and are also required for indispensable cellular functions, such as protein membrane anchoring and signaling (through prenylation) and electron transport (ubiquinone) (17, 18). Essential isoprenoids are derived from two five-carbon building blocks, isopentyl pyrophosphate (IPP) and dimethylallyl pyrophosphate (DMAPP). While mammals, including humans, generate IPP/DMAPP through the mevalonate pathway, *Plasmodium* spp. produce these building blocks through an alternative metabolic route, the methylerythritol phosphate (MEP) pathway (19). The cellular factors that control production of isoprenoid precursors and products in *Plasmodium* spp. are incompletely understood and likely to be highly divergent from those regulatory strategies employed in mammalian cells to control the mevalonate pathway. The essential and parasite-specific MEP pathway thus provides a compelling target for novel antimalarial treatments (20).

The MEP pathway of isoprenoid biosynthesis harnesses two glycolytic intermediates, glyceraldehyde 3-phosphate (G3P) and pyruvate (PYR) which are condensed into 1-deoxy-D-xylulose 5-phosphate (DOXP). This first committed step involves the conversion of DOXP into 2-C-methyl-D-erythritol 4-phosphate (MEP) by the enzyme DOXP reductoisomerase (DXR; E.C. 1.1.1.267). The phosphonate antibiotic fosmidomycin (FSM) is a highly specific, competitive inhibitor of DXR. We have previously employed FSM as a chemical tool to identify novel modes of metabolic regulation in *P. falciparum.* Parasite resistance to FSM is primarily mediated through loss-of-function mutation in one of two closely related small molecule phosphatases, HAD1 (E.C. 3.1.3.23; encoded by PF3D7_1033400) or HAD2 (PF3D7_1226300). HAD1 and HAD2 are members of the large haloacid dehalogenase protein superfamily of phosphotransferases (21,22). Loss of either HAD phosphatase applies metabolic pressure on central carbon metabolism – in particular glycolysis – to divert metabolic precursors toward isoprenoid biosynthesis and achieve FSM resistance. In the case of *had2* mutant parasites, this metabolic dysregulation leads to a fitness cost in asexual replication of the parasite that can be compensated by secondary, hypomorphic alleles in the glycolytic enzyme phosphofructokinase (PFK) (E.C. 2.7.11; PF3D7_0915400) (22). These “second site” mutations in PFK restore FSM sensitivity.

These previous studies highlight a central role for glycolytic regulation in maintaining MEP pathway balance and functionality. Traditionally, PFK serves as the rate-determining step in glycolysis where plants, mammals, and microbes have evolved several layers of regulation. It is therefore not surprising that mutations would arise in PFK to affect glycolytic balance in relation to the MEP pathway. However, not all metabolic states support PFK as the chokepoint in glycolysis. Glycer-aldehyde 3-phosphate dehydrogenase (GAPDH) serves as a critical chokepoint for glycolytic flux under aerobic fermentation, when glucose consumption is high in the presence of oxygen but the absence of TCA cycle respiration (23, 24). This Warburg-like metabolic state is common in rapidly proliferating cells like tumors but also in *Plasmodium* parasites, as glucose nearly acts as the sole energy-producing carbon source with almost no aerobic respiration during the asexual bloodstage. In this state, GAPDH serves as a gate-keeper between upper (energy-consuming “preparatory” phase) and lower (energy-producing “payoff” phase) glycolysis. Beyond its glycolytic function GAPDH functions in multiple other cellular processes including apoptosis, transcription, vesicular trafficking, and heme detoxification (25–30). Currently, the *Plasmodium* GAPDH has not been implicated as a determinant of drug resistance, overall pathogenicity, or a regulator of accessory cellular functions let alone controlling metabolic plasticity despite its primary role as a glycolytic enzyme.

In our continued search of novel modes of FSM resistance we continue to uncover novel biological processes important for parasite function. We present an innovative combination genetic screening strategy that utilizes both enhanced FSM resistance selection and multi-round FSM re-sistance/suppression cycling. Together, these two independent screens converge on GAPDH (E.C. 1.2.1.12; PF3D7_1462800) as an important regulator of glycolytic balance and uncover a secondary accessory role for GAPDH as a heme chaperone that is unique from mammalian GAPDH. This unique mode of metabolic regulation in *Plasmodium* is likely to reveal novel drug targets and therapeutic strategies.

## RESULTS

### Enhanced FSM resistance in malaria parasites selects for GAPDH variant

FSM is a highly specific inhibitor of the MEP pathway, which produces the essential isoprenoid precursors IPP and DMAPP. We have previously shown that *P. falciparum* malaria parasites with loss-of-function mutations in the haloacid dehalogenase, HAD1 (PF3D7_1033400), have elevated levels of MEP pathway metabolites, which confers FSM resistance [half maximal effective concentration (EC_50_) of 4.63 ± 0.46 μM (compared to 0.88 ± 0.05 μM for wild-type (WT) parasites)] (21). To investigate the presence of other metabolic regulators that may be acting in concert with or independently of HAD1, we increased FSM selection pressure in the already resistant *had1* parasite strain (Figure 1). This “enhancer screen” yielded a mixed parasite population that was capable of growth in elevated FSM (10 μM). Two independent clones were derived from this population and likewise exhibited heightened FSM resistance, quantified as >35-fold resistant over WT and >6-fold over their immediate *had1* parent, respectively [33.0 ± 1.02 μM and 31.5 ± 2.76 μM, p<0.0001 compared to WT and *had1* parent with one-way ANOVA and follow-up Bonferroni corrected t-tests (Table 1, Supplemental Figure 1)]. Sanger sequencing verified that this enhanced resistance was not the result of previously described variants in an alternative FSM-resistance locus encoding HAD2 (PF3D7_1226300). Whole genome sequencing on both enhanced FSM resistant clones revealed a novel T626C allele in the gene encoding the canonical glycolytic enzyme, glyceraldehyde 3-phosphate dehydrogenase (GAPDH; PF3D7_1462800). These two clonal FSM resistant strains are hereafter referred to as *had1 gapdh-1a* and *had1 gapdh-1b* (Table 1). The *de novo gadph* allele exists in the second exon of GAPDH and encodes the missense variant GAPDH^I209T^.

**Figure 1:**
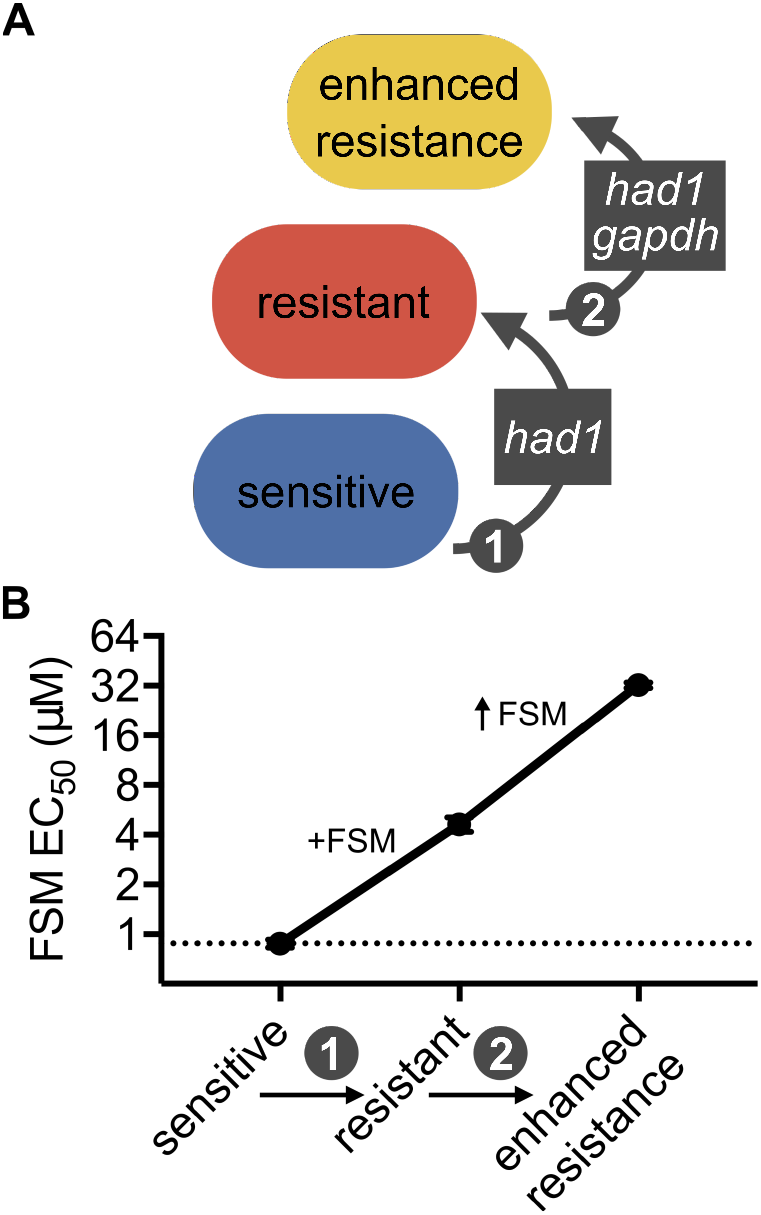
Escalating FSM-resistance screen selects for resistance-associated *gapdh* allele. (**A**) Schematic of the escalating FSM resistance screen and variant alleles identified from whole genome sequencing of FSM-resistant (FSM^R^) *P. falciparum.* (**B**) Screening strategy is shown above the corresponding FSM EC_50_ resistance phenotyping data.

**Table 1.** Summary of strains, genotypes, associated FSM resistance, and NCBI database accession numbers. Strains are grouped according to their separate screening strategies and derived from a 3D7 *Plasmodium falciparum* parental strain. FSM EC_50_ values are determined from three independent biological replicates and displayed as mean ± S.E.M. Genotypes associated with FSM resistance are listed for each strain. Whole genome sequencing data can be found uploaded to NCBI for each strain at the respective Sequence Read Archive (SRA) accession number.

| | Strain | FSM EC<sub>50</sub><br>(μM) | Genotype | Protein Variant | SRA accession<br># |
| --- | --- | --- | --- | --- | --- |
| En-<br>hancer<br>screen | WT | 0.88 ± 0.05 | 3D7 | - | SRR1178414 |
| | *had1* | 4.63 ± 0.46 | *had1*<sup>A28T</sup> | HAD1<sup>K10X</sup> | SRR1178482 |
| | *had1/gapdh-1a* | 33.0 ± 1.02 | *had1*<sup>A28T</sup> *gapdh*<sup>T626C</sup> | HAD1<sup>K10X</sup> GAPDH<sup>I209T</sup> | SRR14929715 |
| | *had1/gapdh-1b* | 31.5 ± 2.76 | *had1*<sup>A28T</sup> *gapdh*<sup>T626C</sup> | HAD1<sup>K10X</sup> GAPDH<sup>I209T</sup> | SRR14929713 |
| Cycling<br>screen | WT | 0.90 ± 0.06 | 3D7 | - | SRR1178414 |
| | *had2* | 4.80 ± 1.20 | *had2*<sup>A469T</sup> | HAD2<sup>R157X</sup> | SRR1745985 |
| | *had2/pfk9* | 0.42 ± 0.04 | *had2*<sup>A469T</sup> *pfk9*<sup>C3617T</sup> | HAD2<sup>R157X</sup> PFK9<sup>T1206I</sup> | SRR14929706 |
| | *had2/pfk9/gapdh-1a* | 1.53 ± 0.36 | *had2*<sup>A469T</sup> *pfk9*<sup>C3617T</sup><br>*gapdh*<sup>T713C</sup> | HAD2<sup>R157X</sup> PFK9<sup>T1206I</sup><br>GAPDH<sup>V238A</sup> | SRR14929705 |
| | *had2/pfk9/gapdh-1b* | 1.72 ± 0.27 | *had2*<sup>A469T</sup> *pfk9*<sup>C3617T</sup><br>*gapdh*<sup>T713C</sup> | HAD2<sup>R157X</sup> PFK9<sup>T1206I</sup><br>GAPDH<sup>V238A</sup> | SRR14929704 |
| | *had2/pfk9/gapdh-2a* | 1.45 ± 0.04 | *had2*<sup>A469T</sup> *pfk9*<sup>C3617T</sup><br>*gapdh*<sup>T713C</sup> | HAD2<sup>R157X</sup> PFK9<sup>T1206I</sup><br>GAPDH<sup>V238A</sup> | SRR14929700 |
| | *had2/pfk9/gapdh-2b* | 1.44 ± 0.08 | *had2*<sup>A469T</sup> *pfk9*<sup>C3617T</sup><br>*gapdh*<sup>T713C</sup> | HAD2<sup>R157X</sup> PFK9<sup>T1206I</sup><br>GAPDH<sup>V238A</sup> | SRR14929699 |
| | *had2/pfk9/gapdh-3a* | 1.60 ± 0.24 | *had2*<sup>A469T</sup> *pfk9*<sup>C3617T</sup><br>*gapdh*<sup>T713C</sup> | HAD2<sup>R157X</sup> PFK9<sup>T1206I</sup><br>GAPDH<sup>V238A</sup> | SRR14929698 |
| | *had2/pfk9/gapdh-3b* | 1.51 ± 0.16 | *had2*<sup>A469T</sup> *pfk9*<sup>C3617T</sup><br>*gapdh*<sup>T713C</sup> | HAD2<sup>R157X</sup> PFK9<sup>T1206I</sup><br>GAPDH<sup>V238A</sup> | SRR15783375 |

### Cycling FSM selection pressure gives rise to separate variant in GAPDH

In parallel to the “enhancer screen” described above, we also employed a second and independent genetic screening strategy (Figure 2). As previously reported, loss of a homologous haloacid dehalogenase, HAD2 (PF3D7_1226300), also results in FSM resistance (EC_50_ = 4.80 ± 1.20 μM), separately from *had1.* Because loss of HAD2 dysregulates central carbon metabolism in *P. falciparum, had2* parasites experience a pronounced fitness cost, such that when FSM selection pressure is removed, the *had2* strain acquires new suppressor mutations in the glycolytic enzyme, phosphofructokinase-9 (PFK9) (22). This *had2 pfk9* strain is again FSM sensitive (EC_50_ = 0.42 ± 0.04 μM) and we have previously determined that PFK9 mutations relieve the dramatic metabolic dysregulation caused by loss of HAD2 (22). To understand the limits of this metabolic plasticity, we iteratively applied FSM selection pressure to the *had2 pfk9* FSM sensitive strain. From three independent selections, we generated FSM resistance and isolated two clones from each selection. All clones exhibited 3-4-fold resistance compared to the *had2 pfk9* strain (Table 1, Supplemental Figure 1). Sanger resequencing confirmed that this resistance was not the result of newly acquired variants in HAD1. To identify the genetic changes responsible for FSM resistance, we performed whole genome sequencing on clones from three independent selections (two clones per selection). Our sequencing converges on a single common locus with a T713C mutation in GAPDH, giving rise to strains *had2 pfk9 gapdh-1a, −1b, −2a, −2b, −3a,* and *-3b,* respectively (Table 1). This new resistant GAPDH allele is also present in the second exon of the gene and encodes the novel missense variant GAPDH^V238A^.

**Figure 2:**
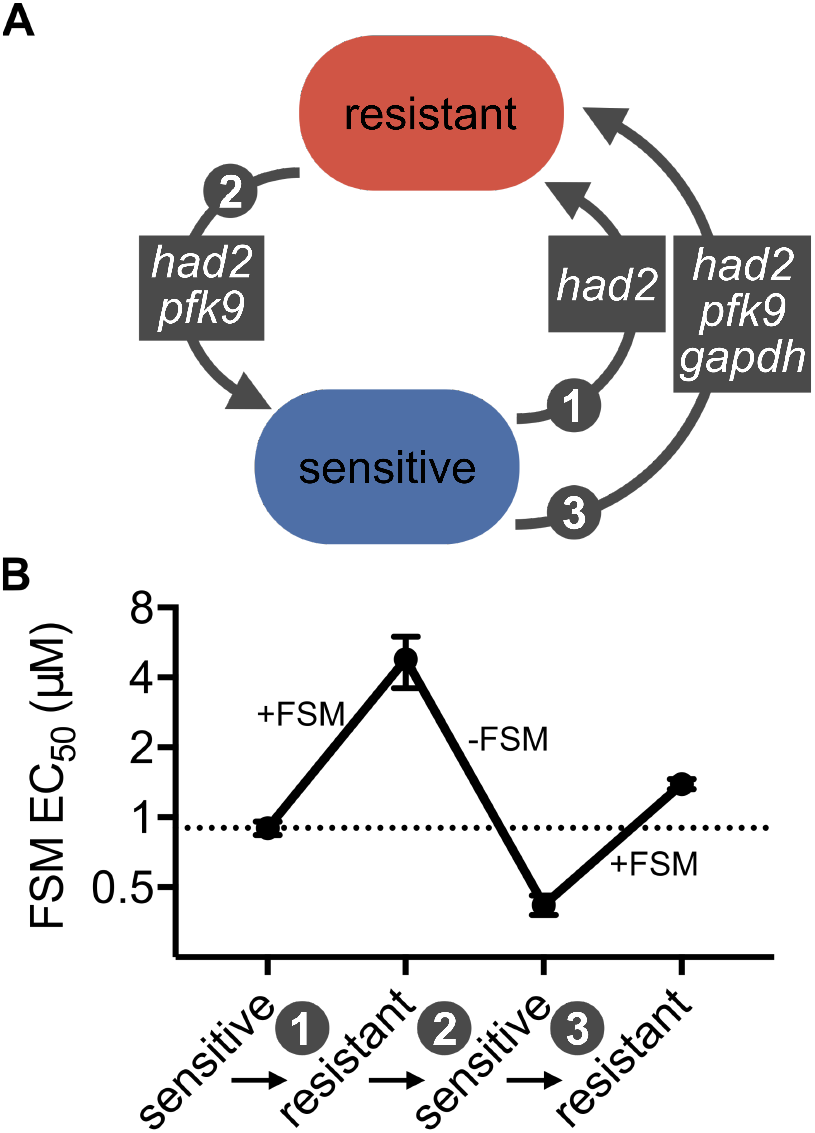
Novel FSM resistance “cycling” screen selects for resistance-associated *gapdh* allele. (**A**) Schematic of the resistance cycling screen and mutant genes identified from whole genome sequencing that yielded FSM^R^ *P. falciparum* parasites from re-sensitized FSM^R^ parasite lines. (**B**) Our screening strategy is shown above the corresponding FSM EC_50_ resistance phenotyping data.

### Two independent selection approaches converge on variants at the dimer interface of GAPDH

Our previous resistance screens unveiled that changes in glycolytic metabolic regulation can be used by malaria parasites to confer altered FSM sensitivity. For this reason, we considered the convergent SNPs in the locus encoding the glycolytic enzyme GAPDH as compelling candidates to mediate FSM resistance. Complete loss of GAPDH function would be unlikely given the essential role for glycolytic function and the need for GAPDH to supply pyruvate for the MEP pathway. To understand how these GAPDH alleles might impact protein function, we modeled the position of the encoded variants on the published tertiary and quaternary structure of *Pf*GAPDH. Both the GAPDH^I209T^ and GAPDH^V238A^ variants are distant from the substrate-binding pocket and are not predicted to directly impact catalysis (Figure 3). Surprisingly, we found that both I209 and V238 are immediately physically adjacent in three-dimensional space. Both residues are present at the base of a relatively disordered S-loop, which is normally stabilized by the oligomerization of the GAPDH homo-tetramer (PDB: 2B4R) (31). Interestingly, we find that not only are I209 and V238 adjacent within a single monomer, they are predicted to directly interact at the hydrophobic interface of two of the homodimer subunits (Figure 3). Classically, hydrophobic surfaces drive proteinprotein interactions. Both FSM-resistant variants alter hydrophobic residues (isoleucine to threonine and valine to alanine, respectively). Therefore, we hypothesized that GAPDH oligomerization was disrupted in our variants and that this was the structural mechanism of FSM resistance in our strains.

**Figure 3:**
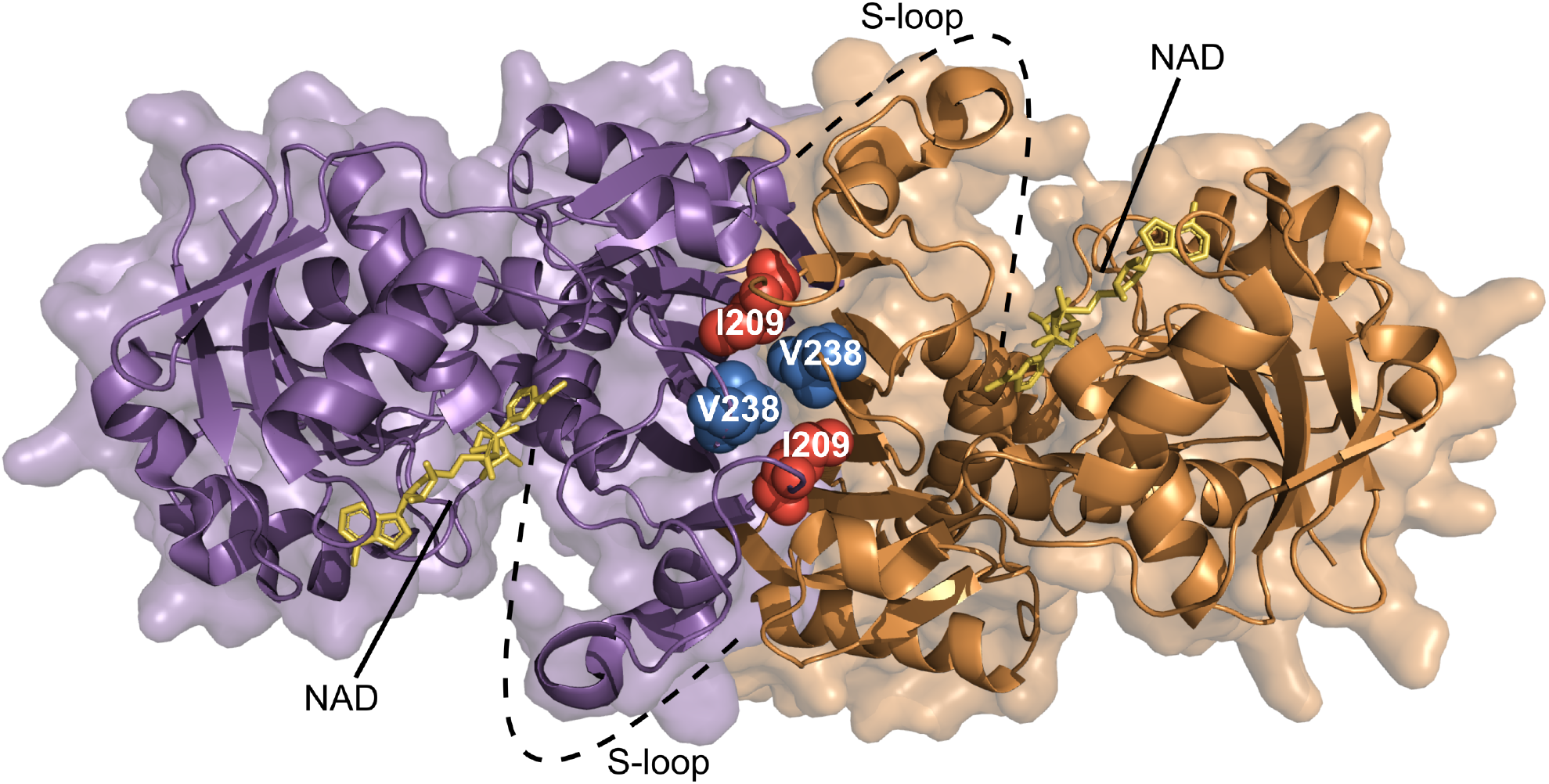
GAPDH mutations from separate FSM resistance screens converge at homodimer interface. Two subunits forming one dimer of the GAPDH tetramer are displayed as ribbon representations in purple and orange with corresponding transparent surfaces. Side chains of the mutation locations are displayed as sphere representations and labeled with white text sitting at the dimer interface and at the base of the S-loop. NAD cofactors are shown as stick figures bound to their respective active sites in yellow (PDB_ID: 2B4R).

### GAPDH tetramer formation is disrupted in FSM-resistant GAPDH variant enzymes

To test this model, we sought to quantify oligomer formation in wild-type GAPDH and its FSM-resistant variants. We performed analytical high-resolution size exclusion chromatography (SEC) (Figure 4), which accurately separates GAPDH monomers and multimers. SEC analysis of GAPDH^WT^ reveals a single dominant peak that corresponds to the tetrameric state (molecular weight of 150 kD, compared to the GAPDH monomer of 37kD). In contrast, both the GAPDH^I209T^ and GAPDH^V238A^ variants exhibit three peaks, which correspond to the tetrameric (150kD), dimeric (83kD), and monomeric (~40kD) states. Loss of the dominant tetramer peak in both FSM-resistant variant proteins confirms that protein-protein interactions in these GAPDH variants are markedly disturbed.

**Figure 4:**
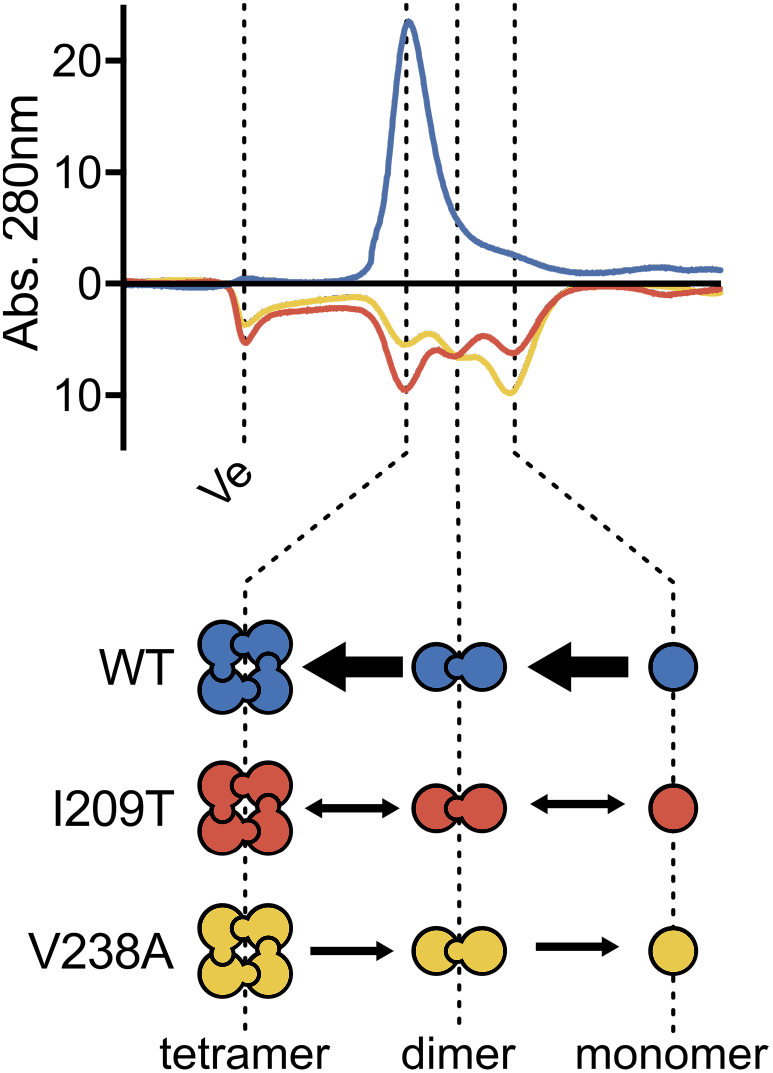
Size exclusion chromatography depicts altered oligomer formation. Size exclu-sion chromatography traces of recombinant proteins with oligomeric states shown below. Ve = void volume of the column. Traces of mutant proteins are inverted below the x-axis.

### Enzymatic characterization of FSM-resistant GAPDH variant enzymes

*Plasmodium* spp. express a phosphorylating GAPDH (E.C. 1.2.1.12) that converts glyceraldehyde 3-phosphate into 1,3-bisphosphoglycerate utilizing inorganic phosphate in an NAD+-de-pendent manner. However, tetramerization of GAPDH is not required for catalysis. Because our FSM-resistant alleles alter residues at the dimer interface that are distant from the active site, we predicted that these GAPDH variants might impact multimerization but would not disrupt the glycolytic function of GAPDH. This model also agrees with our metabolite profiling data, which supports no significant differences in levels of the glycolytic products PYR and LAC. To test this hypothesis, we successfully expressed and purified recombinant wild-type and both FSM-resistant variant proteins (Supplemental Figure 2). Robust glyceraldehyde-3-phosphate dehydrogenase activity was observed for all three protein variants (WT, GAPDH^V238A^, and GAPDH^I209T^), with a decrease in enzyme turnover only detected for GAPDH^V238A^ (*K*cat= 7.0 ± 1.0 s^-1^) but not GAPDH^I209T^ (*K*cat = 55.3 ± 6.1 s^-1^) compared to GAPDH^WT^ (*K*_cat_ = 49.1 ± 7.6 s^-1^), p<0.001 GAPDH^V238A^ vs. GAPDH^WT^ (Figure 5). While modestly reduced enzyme turnover was observed for GAPDH^V238A^ *in vitro,* this was not expected to be biologically meaningful *in vivo,* which agrees with our metabolic profiling data. FSM-resistant and wild-type enzymes shared Michaelis-Menten constants for all three substrates NAD+, GAP, and P_i_ (Supplemental Figure 3).

**Figure 5:**
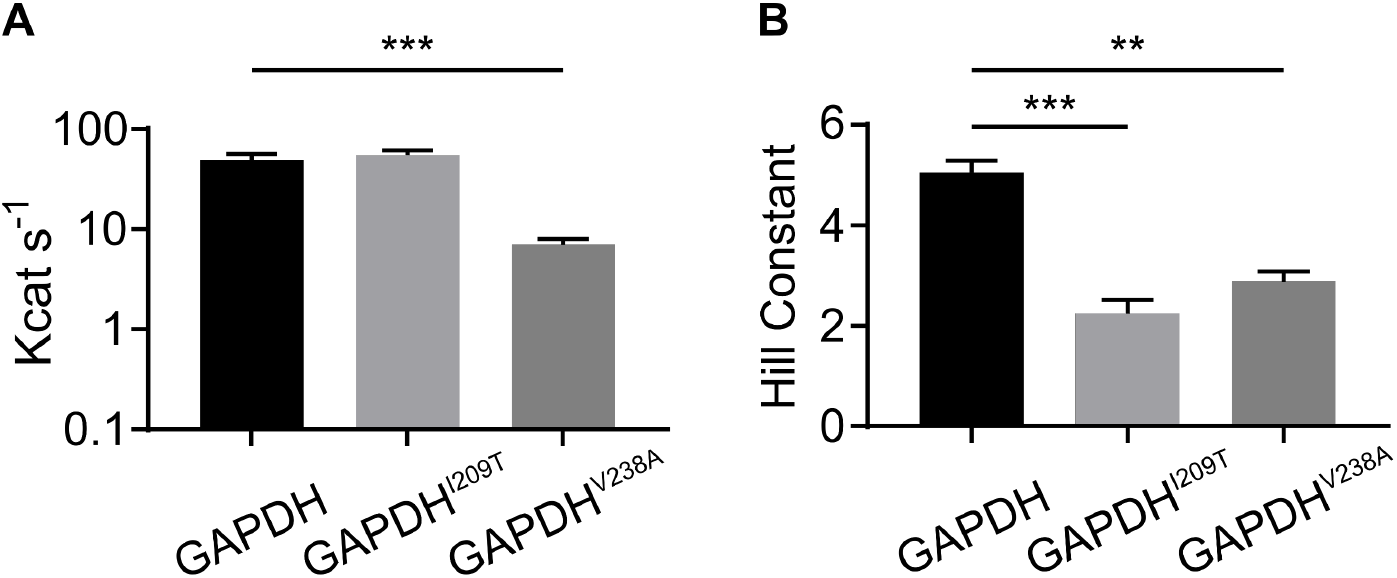
FSM^R^ GAPDH variants support robust catalytic efficiency but altered cooperativity. (**A**) Catalytic efficiency of recombinant GAPDH proteins. A statistically significant decrease in catalytic efficiency is measured for the GAPDH^V238A^ variant compared to WT. (**B**) GAPDH variants display lower cooperativity for phosphate substrate.

In contrast, we find marked differences in inorganic phosphate (Pi) cooperativity between FSM-resistant GAPDH variant enzymes compared to the wild-type enzyme. Cooperativity is a fundamental feature of many multi-subunit proteins, and GAPDH has long been studied as a model of enzymatic cooperativity (32). The binding of small molecule ligands to individual protein subunits leads to conformational changes that induce an increased or decreased ability of substrate to bind to the remaining substrate-binding sites. In the case of GAPDH, P_i_ exhibits positive cooperativity. This cooperativity can be quantified by the Hill constant (H), which approximates the number of substrate binding sites. Both variant enzymes display lower positive cooperativity, as indicated by substantially lower Hill constants (GAPDH^I209T^, H = 2.2 ± 0.; GAPDH^V238A^, H = 2.9 ± 0.2, compared to GAPDH^WT^, H = 5.1 ± 0.2). These findings in conjunction with our size exclusion data suggest a model in which the multimeric assembly is likely disrupted in our fosmidomycin-resistant variant GAPDH enzymes (Figure 5).

### Fosmidomycin-resistant GAPDH variants are also refractory to heme inhibition

GAPDH has been shown to participate in many other cellular processes including apoptosis, transcription, vesicular trafficking, and heme detoxification (25–30). The latter is particularly important with regards to parasite biology (33, 34). The parasite’s voracious appetite for hemoglobin results in large amounts of toxic heme that must be sequestered as hemozoin crystals in the parasite’s food vacuole. Coincidentally, FSM treatment results in disruption in food vacuolar integrity (35). This creates a unique crossroads between GAPDH’s ability to serve as a regulator during metabolic stress, heme toxicity, and drug resistance. While many non-glycolytic roles of GAPDH involve oligomer formation, not all of these non-glycolytic roles are independent from the glycolytic function of GAPDH, especially as a heme chaperone. Therefore, we decided to evaluate whether heme binding was disrupted in our FSM-resistant GAPDH variants. Because heme inhibits the catalytic function of GAPDH, we tested the enzymatic function of our recombinant variant protein compared to wild-type, in the presence or absence of increasing concentrations of free heme. We find that heme inhibits the catalytic function of the wild-type GAPDH protein (EC_50_ = 2.98μM) in a dose-dependent manner. In contrast, FSM-resistant GAPDH variants are highly resistant to heme inhibition (EC_50_ > 10μM, Figure 6). This observation supports the hypothesis that FSM toxicity partly driven by heme toxicity is uncoupled from glycolytic disruption in these GAPDH mutant strains.

**Figure 6.**
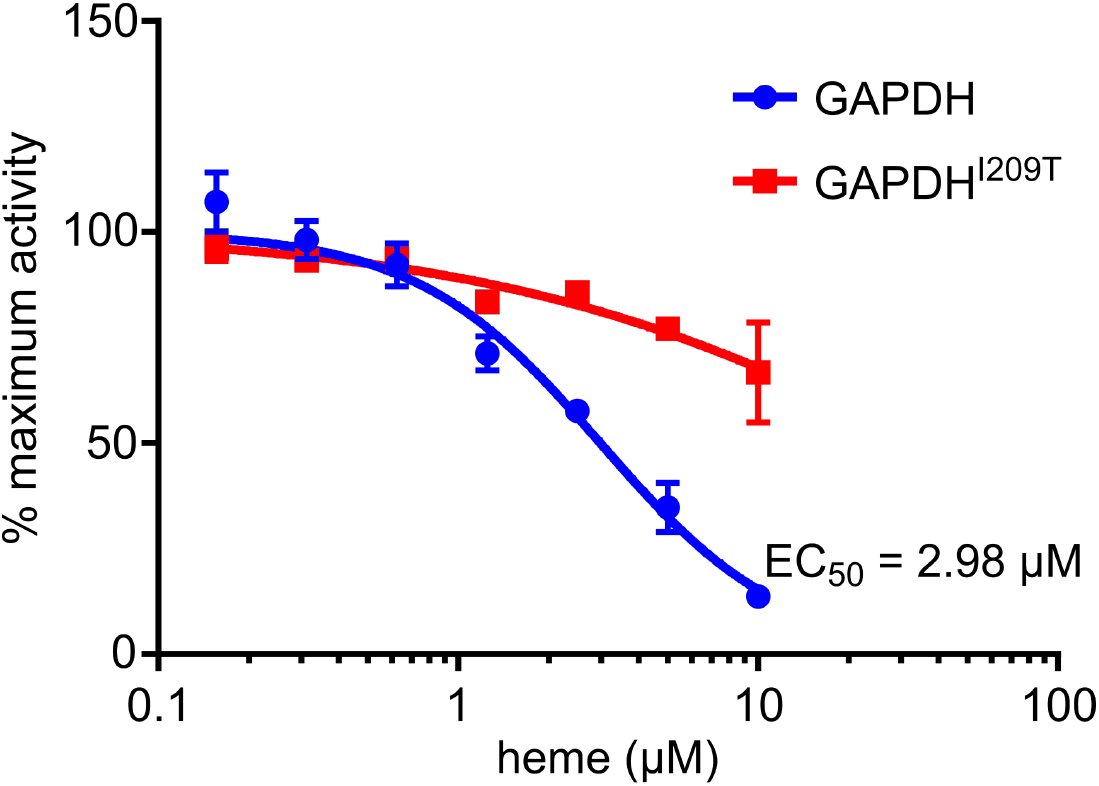
GAPDH mutants are refractory to heme inhibition. Representative EC_50_ of heme against the glycolytic activity of recombinant protein.

### FSM-resistant *gapdh* strains maintain glycolytic output under FSM treatment

To directly test our hypothesis that these GAPDH mutants uncouple the metabolic impacts of FSM, we performed metabolite profiling of MEP pathway and glycolytic intermediates to identify the shared metabolic differences of these *gapdh* strains *(had1/gapdh-a, had2/pfk9/gapdh-2a, – 3a)* compared to their direct parents *(had1, had2/pfk9),* with and without FSM treatment (Figure 7A). A type III two-way ANOVA using a Bonferroni correction for multiple comparisons reveals two metabolites with significant differences: 2-C-methyl-D-erythritol-2,4,-cyclopyrophosphate (MEcPP) with respect to inhibitor treatment (p=0.029), and glycerol 3-phosphate (Gly3P) with respect to genotype (p<0.001) (Figure 7B). As expected, levels of the MEP pathway product MEcPP are significantly reduced for all strains under FSM treatment, which competitively inhibits the MEP pathway upstream of MEcPP production. Although *gapdh* variants have a modest reduction in glycolytic products Lactate (LAC) and Pyruvate (PYR), no significant changes in LAC or PYR levels were observed compared to either immediate parental strain. These data indicate that our FSM-resistance *gapdh* variants nonetheless maintain normal glycolytic output.

**Figure 7:**
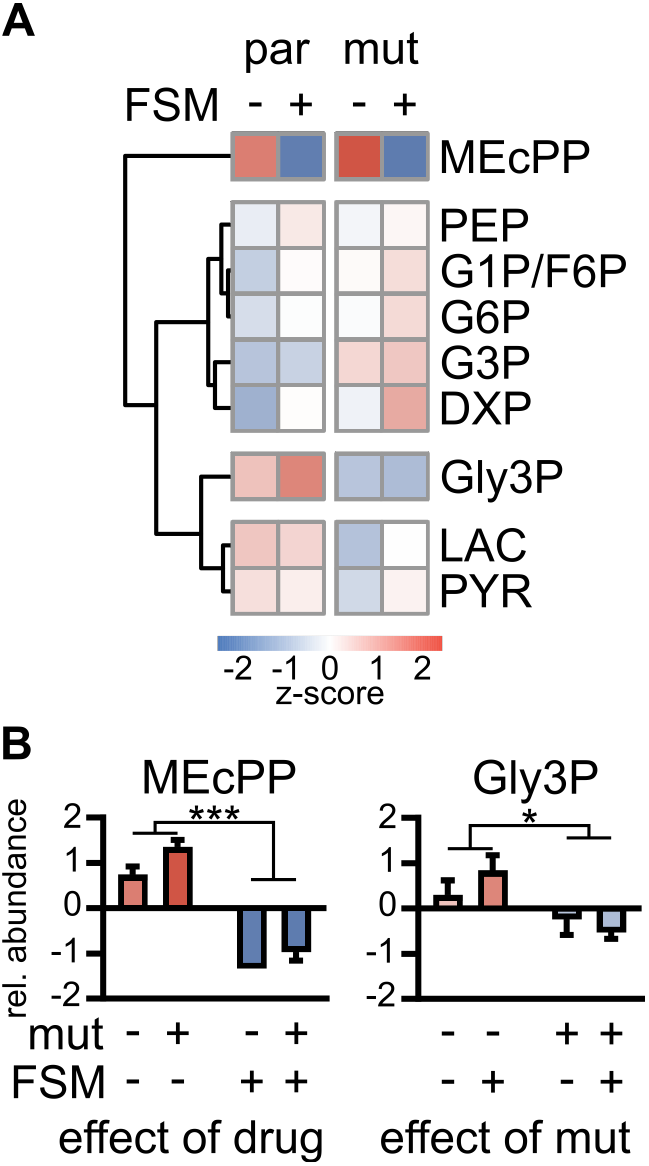
Metabolite profiling indicates healthy glycolytic output. (**A**) Hierarchical clustering of relative abundances for targeted metabolites are displayed comparing parent (par) strains versus alternative *gapdh* alleles (mut) in the presence and absence of FSM treatment. Z-scores represent differences within metabolites across samples. (**B**) Type III two-way ANOVA for targeted metabolites only shows significant differences with respect to drug treatment for 2-C-me-thyl-D-erythritol-2,4,-cyclopyrophosphate (MEcPP) and with respect to genotype for glycerol 3-phosphate (Gly3P).

To determine the role of GAPDH in the context of FSM treatment, we assessed whether GAPDH operates at an inflection point for glycolysis in asexual *P. falciparum* in the presence or absence of FSM treatment. To provide a quantitative measure of these changes, we define “metabolic balance,” as the ratio of the abundance of metabolites upstream of GAPDH [glucose 6-phosphate (G6P), glucose/fructose 1/6-phosphate (G1P/F6P), G3P, and Gly3P] to the abundance of metabolites downstream [phosphoenolpyruvate (PEP), PYR, and LAC] (Figure 8A). We then compared this metabolic balance before and after FSM treatment. We find that FSM dramatically alters metabolic balance in malaria parasites, such that metabolites upstream of GAPDH are elevated relative to metabolites downstream (Figure 8B). In contrast, FSM-resistant *gapdh* mutant strains preserve substrate availability to the lower half of glycolysis (quantified by metabolic balance) in the presence of FSM (p<0.05, unpaired t-test). These data suggest that FSM treatment induces a metabolic disruption in central carbon metabolism that is most pronounced at the level of GAPDH. FSM-resistant GAPDH variants confer FSM resistance through relief of this imbalance (Figure 8C).

**Figure 8:**
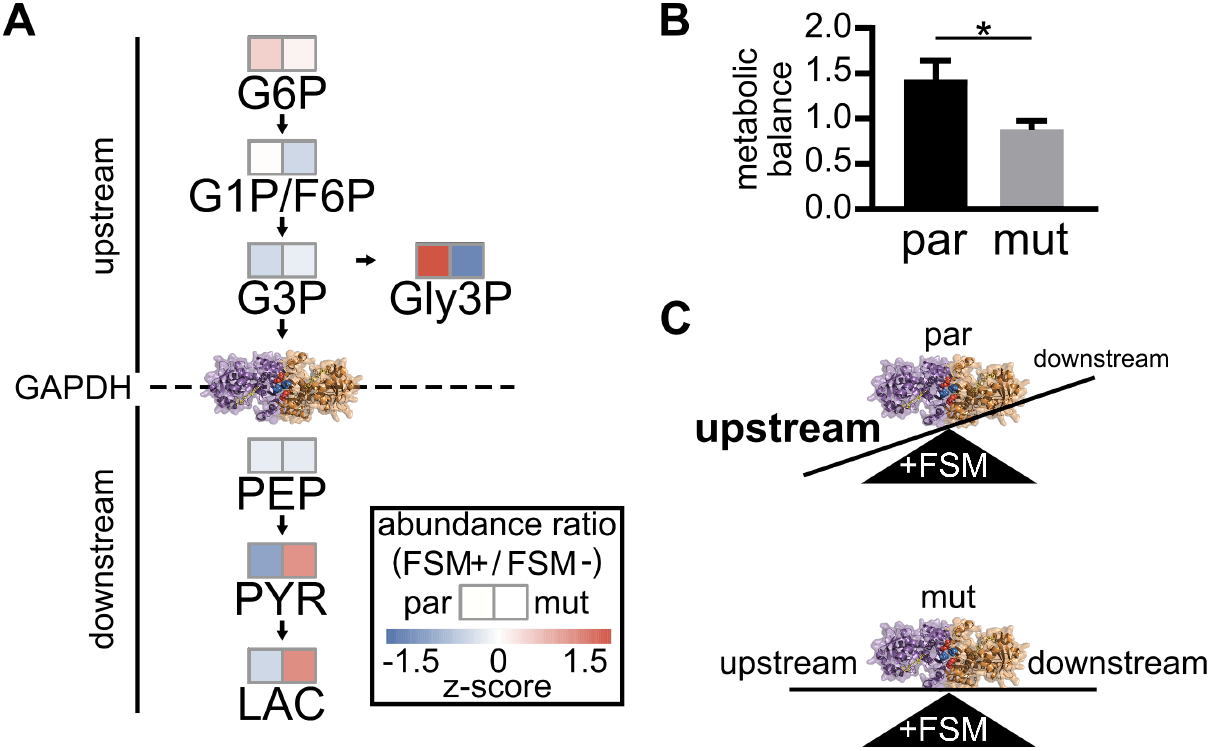
Fosmidomycin-induced metabolic imbalance is resolved in GAPDH mutants. (**A**) Schematic showing metabolites upstream and downstream of GAPDH function. Abundance ratios (FSM + *I* FSM -) are displayed for par and mut strains. Z-scores represent differences within genotypes across metabolites. (**B**) Metabolic balance is displayed as the fold change in total abundance of metabolites upstream over the total abundance of metabolites downstream in the presence of FSM. A t-test shows that FSM induces a significantly different metabolic imbalance in par strains compared to mut strains. (**C**) Schematic illustrating the metabolic imbalance that is resolved by GAPDH mutants under FSM stress.

## DISCUSSION

Two independent FSM drug screening strategies in separate *P. falciparum* strains achieved FSM resistance through a common target, GAPDH. Together, we identified a genetic, molecular, structural, and metabolic mechanism of FSM resistance, which may otherwise not have been identified without these converging screening strategies. Our study reveals an important overall mechanism of metabolic plasticity in the malaria parasite, *Plasmodium falciparum.*

Metabolic plasticity can be described by an organism’s ability to alter metabolism in response to internal and external signals to provide optimal output of metabolic products. Pushing the limits of an organism’s natural metabolic plasticity presents a strategy for identifying metabolic regulators. By targeting an essential downstream metabolic process (isoprenoid production) with a specific chemical inhibitor (FSM), we selected for *Plasmodium falciparum* parasites that have evolved through genetic alterations in metabolic machinery. Thus, FSM resistance continues to serve as an important chemical tool for uncovering essential metabolic regulatory mechanisms in malaria parasites. Resistance to a given inhibitor may be achieved through multiple possible mechanisms, including inhibitor transport (import/efflux), molecular modification, and mutations in the inhibitor target. Each of these possible alternative modes of resistance would provide minimal information about metabolic mechanisms of resistance. Despite these alternative modes of resistance and by pushing the boundaries of FSM’s effects on parasite metabolism, we have identified a new metabolic focal point for achieving FSM resistance.

Given the possible alternative modes of FSM resistance and the continued preference for metabolic alterations in achieving resistance, we have learned an important feature of this small molecule. Our continued selection of metabolic regulatory mechanisms suggests that the fitness cost of metabolic alterations to overcome resistance is far less than the fitness cost of transport or drug target mutations. This observation emphasizes two important points. The first is that the protein target of FSM, DXR, has a high barrier to resistance mutations that render FSM ineffective, as these mutations are not favorable for the essential function of DXR in isoprenoid precursor biosynthesis (36). Our studies also suggest that FSM transport occurs through a route that is likely essential for parasite survival. This observation is counter to what is described in bacteria, where the FSM transporter GlpT is not essential and is readily mutated to achieve FSM resistance (37–39). This stresses the importance of future studies to identify the FSM transporter, likely a novel druggable target.

As a parasite, *Plasmodium falciparum* has a characteristically reduced metabolic robustness given the stable supply of glucose and other nutrients in the blood stream (40). Many of the canonical metabolic modes of regulation that have been described in model organisms do not translate to the parasites’ metabolism, including the lack of an annotated phosphofructose bisphosphatase (FBP) to carry out the reverse step of glycolysis at the level of PFK. Given the reduced metabolic functions within the parasite, much of what remains is a central carbon metabolic skeleton where the metabolites that cannot be scavenged from the host are mostly derived from the glucose uptake of the parasite. Most of this glucose (>90%) flows through glycolysis for ATP production, but a small portion is partitioned toward other essential metabolic processes (41). Our use of FSM as a tool for probing metabolic plasticity has helped us identify the mechanisms that regulate this metabolic partitioning. The current study uncovers a novel mechanism of metabolic regulation through GAPDH.

GAPDH serves an important metabolic role in catalyzing the sixth step of the classical Embden-Meyerhof-Parnas (EMP) pathway of glycolysis, the oxidation of G3P to 1,3-bisphosphoglycerate (1,3BPG), which also yields the reduced product NADH. In glycolysis, GAPDH both follows G3P and proceeds PYR production, which are the two precursor substrates of the MEP pathway. This provides a unique opportunity for GAPDH to maintain balanced substrate flow into the MEP pathway. Beyond its glycolytic metabolic function, GAPDH is involved in multiple other biological processes with roles as diverse as transcriptional regulation, vesicle trafficking, and heme trafficking. Unique to *Plasmodium* spp. is a link between GAPDH’s heme chaperone role and its glycolytic function. The glycolytic function of *Pf*GAPDH is inhibited by heme binding while the mammalian GAPDH is not. This closely ties heme toxicity to metabolic regulation. This is in addition to the regulation of GAPDH function through the formation of tetrameric or dimeric states and the dissociation into is monomeric form. While the ability to perform “alternative” functions may require oligomerization, the glycolytic function does not require oligomer formation.

The most compelling aspect of our discovering novel metabolic control through GAPDH is that two independent screens with the same phenotype of FSM resistance converged on the same genetic locus. Importantly, these mutations were not genetically identical but functionally the same. Our separate non-synonymous mutations generated protein alterations at the same part of the protein responsible for driving multimerization. Disruption of this multimerization changed the ability of heme to inhibit the glycolytic function of GAPDH. Unlike *Hs*GAPDH, *Pf*GAPDH is normally inhibited by heme, as has been previously described. Heme plays a particularly important biological role in parasite biology (34, 42, 43). As the parasite requires hemoglobin digestion for scavenging amino acids it faces a unique problem of detoxifying the heme iron important for the oxygen carrying function of hemoglobin. The parasite mainly achieves this through the polymerization of heme into the characteristic hemozoin crystals found in the parasite food vacuole (42, 44). However, it is still not fully understood how heme is scavenged, transported, and detoxified within the parasite’s cytosol. Recently, GAPDH has been described in other organisms to play a pivotal role in trafficking heme and inserting it into heme requiring apo-proteins, such as iNOS in humans (45, 46). And while a similar function may exist for *Plasmodium* GAPDH, others have hypothesized that GAPDH may serve as a finely tuned heme sensor where the parasite responds to elevated heme by shutting down glycolysis and increasing metabolic flow into the pentose phosphate pathway to generate reducing equivalents (47). This proposed paradigm works well to address oxidative stresses induced by heme only if the cause of rising heme isn’t in conflict with uninhibited GAPDH function. In the case of FSM treatment, where food vacuole integrity is eventually disrupted and therefore presumably causes a concomitant increase in cytosolic heme, the inhibition of GAPDH would further starve the MEP pathway of its precursor metabolites and exacerbate FSM toxicity (35). Our model proposes that under FSM selection pressure parasites respond to reduce this toxicity by eliminating the heme responsiveness of GAPDH in its metabolic function.

As a highly conserved enzyme across the entire kingdom of life, GAPDH has been adapted to perform many other important biological functions. We have shown that *Plasmodium falciparum* is no exception. Roles for GAPDH include regulation of cell death, transcription, oxidative stress, nitrosative stress, vesicle trafficking, and now drug resistance (28, 48, 49). In many of these other possible roles for GAPDH multimerization is likely a factor in performing its function. We now have multimerization mutants in *Plasmodium falciparum* to begin to study these alternative functions. The one-gene one-enzyme hypothesis certainly does not apply to one-function. We aim to understand these possible alternative non-metabolic functions and how they may still play a part in regulating metabolism indirectly.

## METHODS

### Maintaining *P. falciparum* cultures

Parasite strains are derived from 3D7 (MRA-102) as deposited by Daniel J. Carucci at the Malaria Research and Reference Resource Center as part of BEI Resources and established by the National Institutes of Allergy and Infectious Diseases. Parasite cultures were maintained in a suspension of human erythrocytes at 2% hematocrit in complete media (RPMI-1640, Millipore-Sigma, supplemented with 27mM sodium biocarbonate, 11mM glucose, 5mM HEPES, 1mM sodium pyruvate, 0.37mM hypoxanthine, 0.01 mM thymidine, 10 μg ml^-1^ gentamycin, and 0.5% Albumax, Life Technologies) under 5% O2/5% CO2/90% N_2_ atmosphere at 37°C. Human erythrocytes were obtained from banked blood kindly provided by the Saint Louis Children’s Hospital. Donated blood is filtered to remove leukocytes. All blood is triple washed using incomplete media (RPMI-1640, Millipore-Sigma) and stored at 50% hematocrit at 4°C. Erythrocytes are used up to 1 month past the clinical expiration date.

### Generating enhanced FSM resistant *P. falciparum*

Strain E1 was created as previously published and described (21). As part of a larger enhanced resistance screening strategy strain E1 was first cloned by limiting dilution. The newly obtained cloned was deemed strain E1-C12. From this clone independent selections were considered as separated wells. Parasites were continually cultured as described with the addition of 10 μM FSM. Cultures that were positive for parasite growth were isolated and verified for FSM resistance as compared to strain E1-C12. Enhanced FSM-resistant strain 59 was cloned by limiting dilution. Two clones 59a and 59b were isolated, confirmed for FSM resistance, and submitted for whole genome sequencing.

### Generating FSM resistance strains in FSM resistance suppressed *P. falciparum*

Strains E2 and E2-S1 were created as previously published and described (21,22). Strain E2-S1 was first cloned by limiting dilution. The newly obtained clone was deemed strain E5. From this clone independent selections were performed in separated wells. Parasites were continually cultured as described with the addition of 0.5 μM FSM. Cultures that were positive for parasite growth were isolated and verified for FSM resistance as compared to strain E5. Four independent FSM resistant isolates were selected and named E5-1, E5-2, E5-3, and E5-4. Each isolate was cloned by limiting dilution and two subclones were isolated for each, confirmed for FSM resistance, and submitted for whole genome sequencing.

### Determination of half maximal FSM growth inhibition

Parasite cultures were diluted to 1% parasitemia and delivered to a 96-well plate along with an uninfected erythrocyte control. Serial dilutions of FSM were delivered along with a solvent control and cultures were incubated under normal culture conditions for three days. DNA was quantified using Picogreen (Life Technologies), as previously described (50). Half-maximal effective concentration (EC_50_) from nonlinear regression of normalized background-subtracted measurements of at least three biological replicates (GraphPad Prism).

### Whole genome sequencing and variant discovery

Isolated DNA was submitted to the Washington University Genome Technology Access Center for library preparation, sequencing, read alignment and variant analyses. One microgram of genomic DNA was sheared, end repaired, and adapter ligated. Sequencing was performed on an Illumina HiSeq 2500 in Rapid Run mode to generate 101-bp paired-end reads. Demultiplexed reads were aligned to the *Plasmodium falciparum* 3D7 reference genome (PlasmoDB v7.2) using the short-read aligner Novoalign. Variants were called using samtools with a quality score greater than 37 and a minimum read depth of 5. Variants were annotated using snpEff. Variants present in the wild-type strain were removed from further analysis. Sequencing reads are available in the Sequence Read Archive (SRA) of the National Center for Biotechnology Information (NCBI), part of BioProject PRJNA222697. Individual SRA Accession numbers can be found in Table 1.

### Sanger sequence verification of *PfGAPDH*

The regions with the gene that contained mutations were amplified from genomic DNA isolated from each parasite strain using the following primers: 5’-ATGGCAGTAACAAAACTTGG-3’ and 5’-TTAGTTGTTAGTAATGTGTACGG-3’. Purified PCR products were sequenced by the Protein and Nucleic Acid Laboratory at Washington University using BigDye Terminator v3.1 Cycle Sequencing reagents (Life Technologies) using primers: 5’-CCACCAATATGATACCAAAC-3’, 5’-TCAGTGTATCCTAAGATTCC-3’. Sanger sequencing chromatograms were aligned to *gapdh* (PF3D7_1462800) using the SeqManPro software (DNAStar).

### Recombinant expression, purification, and size exclusion chromatography of *PfGAPDH*

An *E. coli* codon-optimized *Pf*GAPDH was produced (Genewiz) with N-term and C-term modifications for improved solubility as previously described (31). Mutations for our corresponding mutant proteins were introduced using site-directed mutagenesis. Each sequence was then inserted via ligation-independent cloning into the isopropyl β-D-1-thiogalactopyranoside (IPTG) inducible BG1861 expression vector. This creates an N-terminal 6x-His tag fusion protein used for nickel purification. Expression plasmids were transformed into One Shot BL21(DE3) *E. coli* cells (Thermo Scientific). Overnight starter cultures were diluted 1:1000 and grown to an optical density (O.D.) of ~0.6 where 1mM IPTG was added for 2 hours at 37°C. Cells were spun and stored at −80°C.

Expressed proteins were purified from cells using a sonication lysis buffer containing 1 mg/ml lysozyme, 20mM imidzazole, 1mM dithiothreitol, 1mM MgCl_2_, 10mM Tris HCl (pH7.5), 30 U benzonase (EMD Millipore), 1mM phenylmethylsulfonyl fluoride (PMSF), and EDTA-free protease inhibitor tablets. Lysates were clarified using centrifugation and proteins were purified via nickel agarose beads (Gold Biotechnology), eluted with 300mM imidazole, 20mM Tris-HCl (pH 7.5) and 150mM NaCl. Eluted proteins were further purified via size exclusion chromatography using a HiLoad 16/60 Superdex 200 gel filtration column (GE Healthsciences) using an AKTAExplorer 100 FPLC (GE Healthsciences). Fast protein liquid chromatography buffer contained 100mM T ris-HCl (pH 7.5), 1mM MgCl_2_, 1mM DTT, and 10% w/v glycerol. Fractions containing purified protein were pooled, concentrated to ~2mg/ml as determined via Pierce BCA Protein Assay Kit (ThermoFisher), and stored by adding 50% glycerol for storage at −20°C.

### Determination of *PfGAPDH* enzyme activity

Enzyme activity was determined spectrophotometrically at 340nm. The reaction was measured in buffer containing 10mM Tris-HCl pH 7.5, 200mM sodium chloride, 1mM magnesium chloride, 1mM DTT, 0.1 mg/mL bovine serum albumin (BSA), 25mM sodium phosphate (NaP_i_), and 5mM NAD+. Recombinantly expressed proteins were added and allowed to incubate at 37°C for 10 minutes before starting the reaction with D-glyceraldehyde 3-phosphate (G3P). Assays were performed in a 96-well plate format and read by a microplate spectrophotometer (BMG Labtech). Conversion of NAD+ to NADH was determined using the extinction coefficient of 6,220 M^-1^cm^-1^.

### Michaelis-Menten kinetics

100μM NAD^+^, 64μM G3P, and 500μM P_i_ were each serial diluted according to the reaction mix described above. The activity assays were performed as described above. Nonlinear regression of the slopes of the different substrate concentrations yielded a Michaelis-Menten curve with corresponding K_m_, and V_max_ values for substrates G3P and NAD+. Allosteric sigmoidal plots were fitted for determining V_max_, K_half_, and Hill constants for P_i_. (GraphPad Prism).

### Heme inhibition of *Pf*GAPDH

*A* fresh stock of hemin (2.5mM) was prepared by dissolving hemin in 100mM NaOH with 5% DMSO. The stock hemin solution was diluted further to 1mM in 100mM NaOH with 5% DMSO. Hemin was diluted 1:5 in GAPDH reaction buffer as described above. Serial dilutions of hemin were added to recombinant enzyme and allowed to incubate for 10 minutes at room temperature before being added to the standard GAPDH assay reactions described above.

### Sample collection for LC/MS metabolite quantitation

*Plasmodium falciparum* strains were split into independent cultures at least two weeks before sample collection. Cultures were grown to ~10% parasitemia in 30mL volumes with 4% hematocrit. Cultures were then synchronized with 5% sorbitol treatment in successive rounds until >75% of parasites were in ring-stage growth. Ring-stage parasites were treated ± FSM (5μM) (Life Technologies) for 12 hours. Parasite pellets were collected via erythrocyte lysis induced with 0.1% saponin and centrifugation. Pellets were washed with PBS and flash frozen with LN2 for storage at-80°C.

### LC/MS sample extraction and analysis

MEP, TCA, glycolysis and PP pathway intermediates were extracted via the addition of glass beads (212-300 u) and 600 μL chilled H_2_O: chloroform: methanol (3:5:12 v/v) spiked with PIPES (piperazine-N,N-bis(2-ethanesulfonic acid) as internal standard. The cells were disrupted with the TissueLyser II instrument (Qiagen) using a microcentrifuge tubes adaptor set pre-chilled for 2 min at 20 Hz. The samples were then centrifuged at 16,000 g at 4 °C, the supernatants collected, and pellet extraction repeated once more. The supernatants were pooled and 300 μL chloroform and 450 μL of chilled water were added to the supernatants. The tubes were vortexed and centrifuged. The upper layer was transferred to a new tube and dried using a speed-vac. The pellets were redissolved in 100 μL of 50% acetonitrile.

For LC separation of the MEP intermediates, a Luna-NH2 column (3 um, 150 x 2 mm, Phenomenex) was used flowing at 0.4 mL/min. The gradient of the mobile phases A (20 mM ammonium acetate, pH 9.8, 5% ACN) and B (100% acetonitrile) was as follows: 60% B for 1 min, to 6% B in 3 min, hold at 6% B for 5 min, then back to 60% B in 0.5 min. For LC separation of the TCA/Glycolysis/PPP intermediates, an InfinityLab Poroshell 120 HILIC (2.7 um, 150 x 2.1 mm, Agilent) was used flowing at 0.5 mL/min. The gradient of the mobile phases A (20 mM ammonium acetate, pH 9.8, 5% ACN) and B (100% acetonitrile) was as follows: 85% B for 1 min, to 40% B in 9 min, hold at 40% B for 2 min, then back to 85% B in 0.5 min. The LC system was interfaced with a Sciex QTRAP 6500+ mass spectrometer equipped with a TurboIonSpray (TIS) electrospray ion source. Analyst software (version 1.6.3) was used to control sample acquisition and data analysis. The QTRAP 6500+ mass spectrometer was tuned and calibrated according to the manufacturer’s recommendations. Metabolites were detected using MRM transitions that were previously optimized using standards. The instrument was set-up to acquire in negative mode. For quantification, an external standard curve was prepared using a series of standard samples containing different concentrations of metabolites and fixed concentration of the internal standard. The limit of detection for deoxyxylulose 5-phosphate (DOXP), methylerythritol phosphate (MEP), cytidine diphosphate methylerythritol (CDP-ME), and methylerythritol cyclodiphosphate (MEcPP) was 0.0064 μM for a 10 μL injection volume. The limit of detection for TCA, Glycolysis and PPP intermediates were as follows: KGLU 0.01 μM; ACO 0.15 μM; MAL and SUC 0.2 μM; FUM and LAC 1 μM; GLU and PYR 2 μM; G1P/F6P and G6P 0.5 μM; GADP 1 μM; 2PGA/3PGA and PEP 2 μM; Gly3P 0.25 μM; Ru5P 1 μM, S7P 2 μM.

### Metabolite data processing and analysis

Metabolite abundances were log-transformed with pareto scaling to generate normally distributed data. Metabolites with abundances below the limit of detection were imputed at half the value of the lowest measured abundance for that metabolite. Samples were collapsed into two groups, those with parent genotypes and those with mutations in GAPDH. A type III two-way analysis of variance (ANOVA) was performed on each metabolite to test for significant differences across genotype, drug treatment, and/or interaction. A Bonferroni correction was applied to adjust for multiple comparisons. All analysis were performed using the R package of MetaboAnalyst.

## ACKNOWLEDGMENTS

A.J.J. was supported by NIH T32GM007067 and 2T32AI007511. A.M.G. was supported by the Monstanto Excellence Fund Graduate Fellowship. L.K.M. was supported by the Mr. and Mrs. Spencer T. Olin Fellowship for Women. A.O.J. is supported by NIH/NIAID R01-AI103280, R21-AI123808, and R21-AI130584, and AOJ is an Investigator in the Pathogenesis of Infectious Diseases (PATH) of the Burroughs Wellcome Fund.

We thank the Genome Technology Access Center at the McDonnell Genome Institute at Washington University School of Medicine for help with genomic analysis. The Center is partially supported by NCI Cancer Center Support Grant #P30 CA91842 to the Siteman Cancer Center and by ICTS/CTSA Grant# UL1TR002345 from the National Center for Research Resources (NCRR), a component of the National Institutes of Health (NIH), and NIH Roadmap for Medical Research. This publication is solely the responsibility of the authors and does not necessarily represent the official view of NCRR or NIH.

---

**Supplemental Figure 1.**
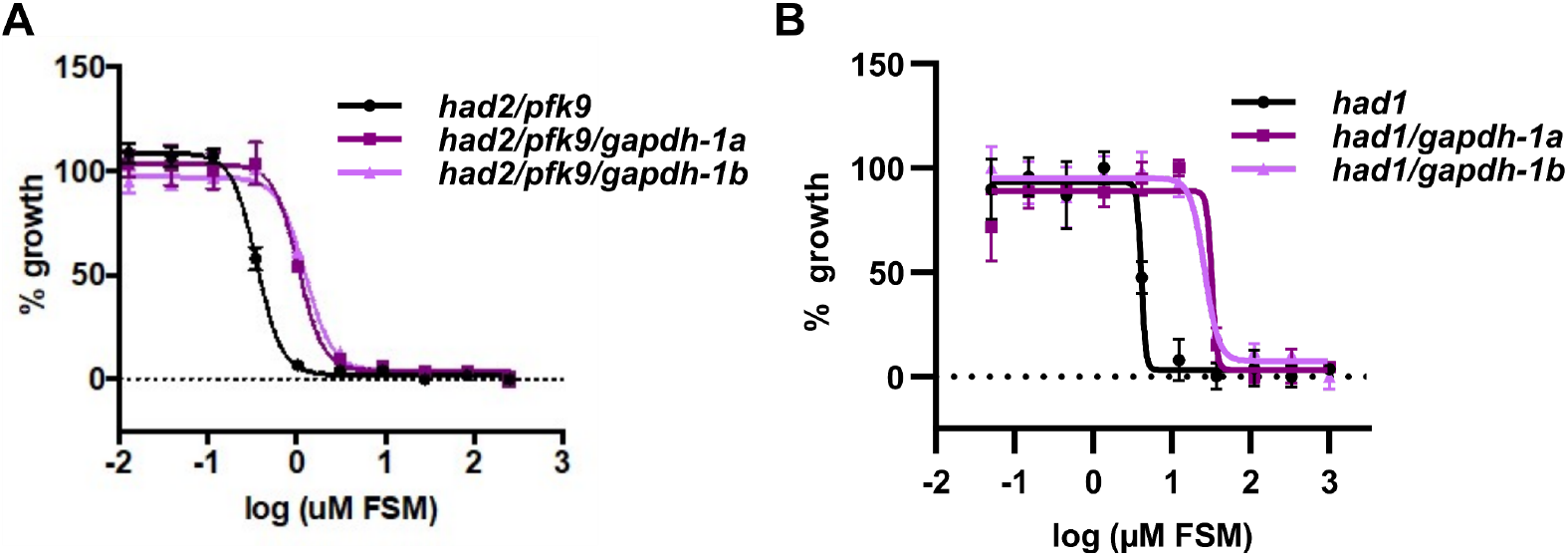
Representative EC_50_ curves of enhancer and cycling screen strains.

**Supplemental Figure 2.**
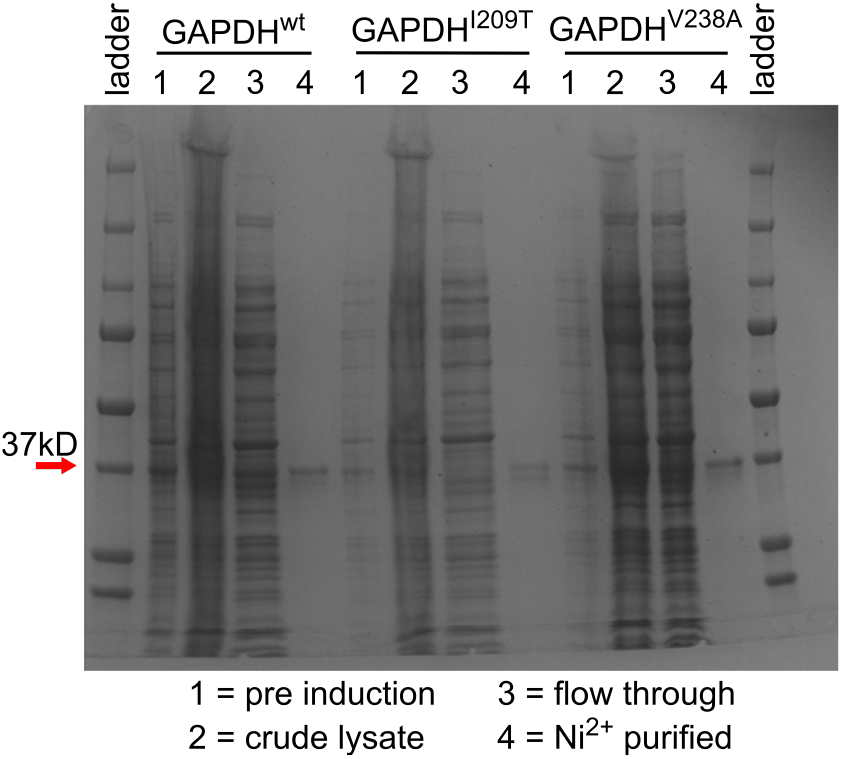
Recombinant protein Coomassie gel.

**Supplemental Figure 3.**
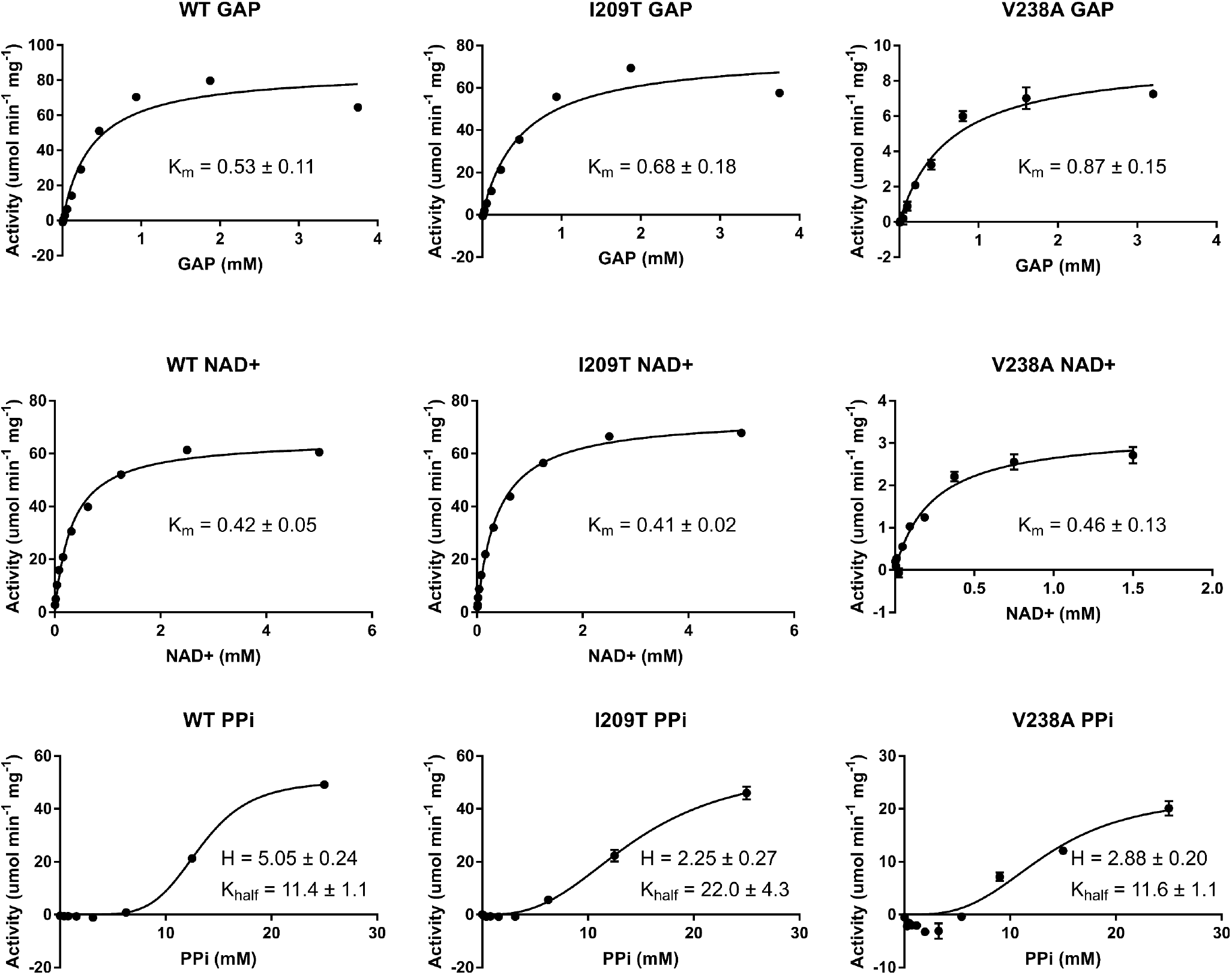
Michaelis-Menten Kinetics.

**Supplemental Table. Whole genome sequencing results.**

